# Revealing biases in the sampling of ecological interaction networks

**DOI:** 10.1101/328245

**Authors:** Marcus A. M. de Aguiar, Erica A. Newman, Mathias M. Pires, Justin D. Yeakel, David H. Hembry, Carl Boettiger, Laura A. Burkle, Dominique Gravel, Paulo R. Guimarães, James L. O’Donnell, Timothée Poisot, Marie-Josée Fortin

**Affiliations:** Universidade Estadual de Campinas, Unicamp, Campinas, São Paulo, Brazil; USDA Forest Service, Pacific Wildland Fire Sciences Lab, Seattle, WA, USA; School of Natural Resources and the Environment, University of Arizona, Tucson, AZ, USA; School of Natural Sciences, University of California, Merced, CA, USA; Santa Fe Institute, Santa Fe, NM, USA; Department of Ecology and Evolutionary Biology, University of Arizona, Tucson, AZ, USA; Department of Environmental Science, Policy, and Management, University of California, Berkeley, CA, USA; Montana State University, Department of Ecology, Bozeman, MT, USA; Département de Biologie, Université de Sherbrooke, Sherbrooke, QC, Canada; Departamento de Ecologia, Instituto de Biociências, Universidade de São Paulo, São Paulo, SP, Brazil; School of Marine and Environmental Affairs, University of Washington, Seattle, WA, USA; Québec Centre for Biodiversity Sciences, Montréal, QC, Canada; Département de Sciences Biologiques, Université de Montréal, Montréal, QC, Canada; Department of Ecology & Evolutionary Biology, University of Toronto, Toronto, ON, Canada

## Abstract

The structure of ecological interactions is commonly understood through analyses of interaction networks. However, these analyses may be sensitive to sampling biases in both the interactors (the nodes of the network) and interactions (the links between nodes), because the detectability of species and their interactions is highly heterogeneous. These issues may affect the accuracy of empirically constructed ecological networks. Yet statistical biases introduced by sampling error are difficult to quantify in the absence of full knowledge of the underlying ecological network’s structure. To explore properties of large-scale modular networks, we developed *EcoNetGen*, which constructs and samples networks with predetermined topologies. These networks may represent a wide variety of communities that vary in size and types of ecological interactions. We sampled these networks with different sampling designs that may be employed in field observations. The observed networks generated by each sampling process were then analyzed with respect to the number of components, size of components and other network metrics. We show that the sampling effort needed to estimate underlying network properties accurately depends both on the sampling design and on the underlying network topology. In particular, networks with random or scale-free modules require more complete sampling to reveal their structure, compared to networks whose modules are nested or bipartite. Overall, the modules with nested structure were the easiest to detect, regardless of sampling design. Sampling according to species degree (number of interactions) was consistently found to be the most accurate strategy to estimate network structure. Conversely, sampling according to module (representing different interaction types or taxa) results in a rather complete view of certain modules, but fails to provide a complete picture of the underlying network. We recommend that these findings be incorporated into field sampling design of projects aiming to characterize large species interactions networks to reduce sampling biases.

**Author Summary:** Ecological interactions are commonly modeled as interaction networks. Analyses of such networks may be sensitive to sampling biases and detection issues in both the interactors and interactions (nodes and links). Yet, statistical biases introduced by sampling error are difficult to quantify in the absence of full knowledge of the underlying network’s structure. For insight into ecological networks, we developed software *EcoNetGen* (available in R and Python). These allow the generation and sampling of several types of large-scale modular networks with predetermined topologies, representing a wide variety of communities and types of ecological interactions. Networks can be sampled according to designs employed in field observations. We demonstrate, through first uses of this software, that underlying network topology interacts strongly with empirical sampling design, and that constructing empirical networks by starting with highly connected species may be the give the best representation of the underlying network.

## Introduction

Network theory provides an efficient way to represent and characterize the structure of ecological systems by organizing the complex relationships between species as graphs, where nodes represent the species, and links represent their interactions (Pascual & Dunne 2005). Field observations used to construct species interaction networks can be effort-intensive, so empirical networks often focus on a single module, that is, a subset of highly interconnected species. Furthermore, a field ecologist may attempt to exhaustively sample the species interacting in a delimited area, while ignoring interactions and species that occur outside that area. This stems from a fundamental challenge in community ecology as a whole: establishing the boundaries of the system of interest (Morin 2009).

Because empirical networks are often constructed with a focus on a given type of interaction and by sampling interactions of a particular taxonomic group within a locality (Hall & Raffaelli 1993; Bascompte & Jordano 2007), the largest empirical ecological interaction networks typically include no more than a few hundred species; often many fewer. Nonetheless, these represent sub-networks within a more complete ecological network, which will include many more species interacting in multiple qualitatively different ways (Box 1) (Fontaine et al. 2011; Pilosof et al. 2017). For example, a plant-pollinator network focusing on insects is one module of a larger network that includes pollinators from other taxonomic groups, as well as the consumers of these species and their parasites, and so on. The study of empirical networks relies on the reasonable assumption that the ecological and evolutionary dynamics of each subnetwork can usually be investigated independently (Lewinsohn et al. 2006). Yet there are situations in which neglecting the effects of other interactions and species outside the delineated boundaries may lead to incomplete or incorrect conclusions (Ings et al. 2009; Fontaine et al. 2011; Mello et al. 2011a, 2011b; Rivera-Hutinel et al. 2012). The increased interest in larger ecological networks encompassing several groups and types of networks, such as multilayered networks (Pilosof et al. 2017), and development of tools for their study makes this problem especially timely for ecology.

### Box 1. Variations in structure among ecological interaction networks

Network topology can vary greatly from one part of the network to another, and influence the conclusions drawn about the underlying network. For example, interactions among certain groups of species form subnetworks characterized by high degrees of modularity and reciprocal specialization, as is the case with some ant-myrmecophyte networks (Guimarães et al. 2007), clownfish-anemone networks (Fautin & Allen 1997; Ollerton et al. 2007), and other networks where interactions are symbiotic (Hembry et al. 2018). Conversely, mutualisms such as those between plants and their pollinators or seed dispersers (Bascompte et al. 2003), or the interactions between generalized predators or herbivores with the resources they consume (Pires and Guimarães 2013) are highly nested, where the interactions of specialists are a nested subset within those of generalists. Yet, as we look at broader scales that include multiple habitats, taxonomic groups, and/or interaction types, a modular organization tends to emerge (Baskerville et al. 2011; Olesen et al. 2007; Donatti et al. 2011), with each module having unique structural properties (Lewinsohn et al. 2006; Fontaine et al. 2011). More complete ecological networks may emerge from the aggregation of multiple types of interactions, as well as the various and sometime unique structures such interactions form, and can be represented as large, modular networks. However, it is unknown whether and to what extent different sampling strategies might bias our understanding of the underlying network structure (e.g. Jordano 2016; Fründ et al. 2016; Vizentin-Bugoni et al. 2016).

**Table:**
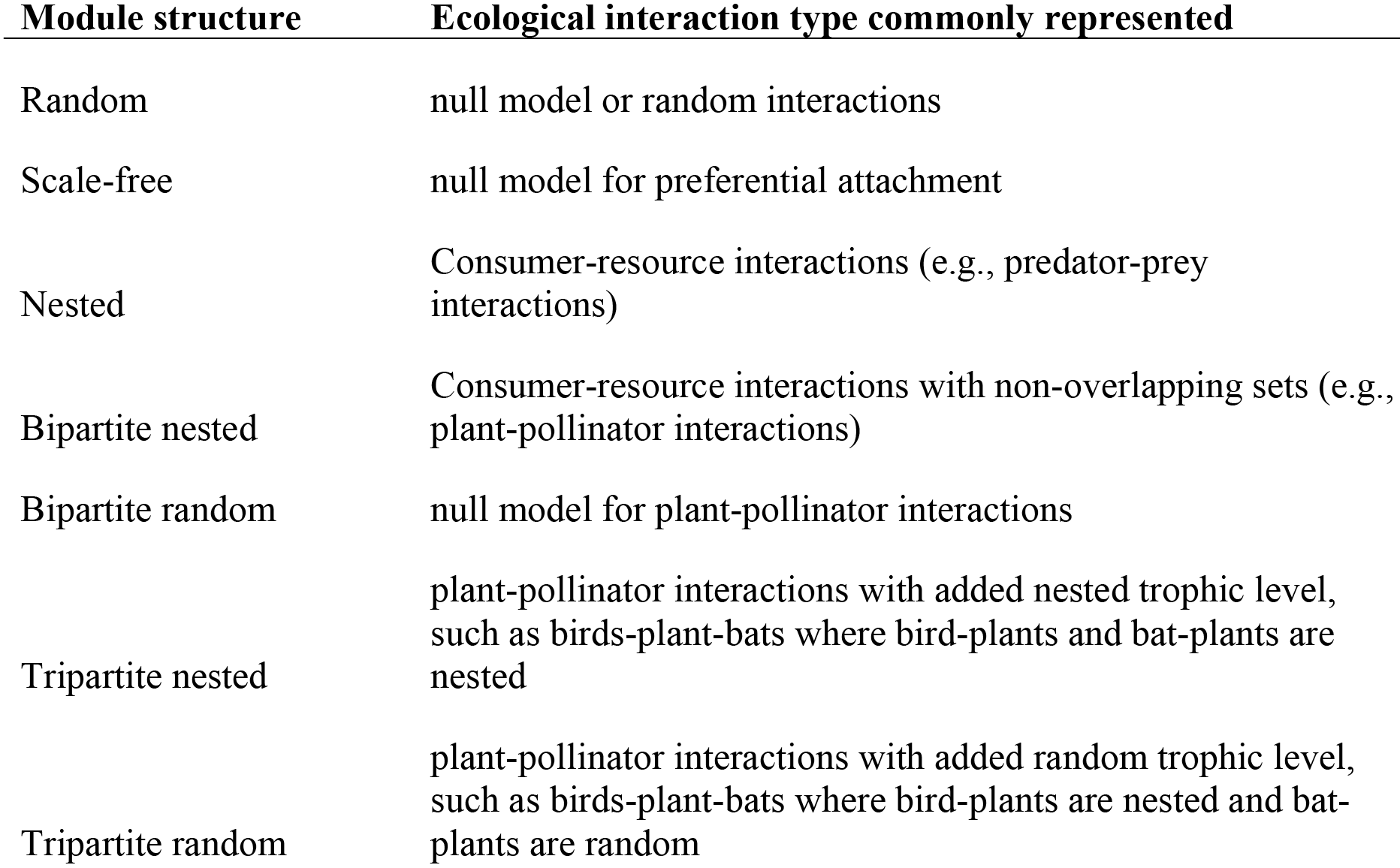
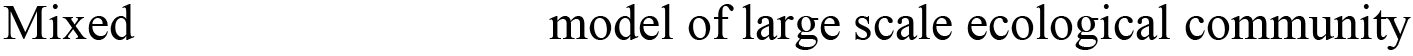
Simulated modules represent certain types of ecological interactions, each of which are known to have different associated structural characteristics.

The question is then: how much of the underlying, complete ecological network can be observed by sampling a subset of its species (nodes) and their associated interactions (links)? The effectiveness of field sampling in capturing the underlying complete network may depend on (1) the underlying topology of the complete network, (2) the sampling technique itself, and (3) the potential interplay between the network topology and the sampling strategy.

We investigated the above questions through a new software, *EcoNetGen*, developed initially for this project in Python using Fortran, and now available in the R programming language (de Aguiar et al. 2018) and on CRAN. *EcoNetGen* contains the scripts *NetGen*, which generates interaction networks with predetermined properties (including structure of the modules within the network, as well as the frequency of modules with particular structures), and *NetSampler*, which samples sets of nodes from the full network according to a chosen technique, and then compare the observed network against the full, complete network. Specifically, we examined how the interplay between network structure and sampling design alters our inference on the among-module connectedness in networks, and draw conclusions about the sampling design that captures the most accurate picture of the complete network for a given topology. This allowed us to evaluate how different sampling strategies often used in the field might alter observational accuracy, identify whether specific sampling designs produce more reliable estimates of the underlying network structure, and to what extent such designs can be confounded or enhanced by alternative arrangements of the underlying species interactions.

To better understand the factors that may affect the match between the topology of the sampled and underlying networks, we focused on simulated networks in which we can control the initial structure and properties such as size and connectance. We also focused on simplified modules that depict well-defined structures instead of trying to encompass all the variability that can be found in nature. The investigations presented here are theoretical investigations of a range of possible network topologies, rather than a comparison to field-sampled data, as any empirical network available for analyses is already a sampled version and contains the effects of sampling biases. By building upon simplified structures we have a greater control on what is affecting network properties. We expect that the range of network topologies illustrated here can adequately describe many empirically observed patterns and give insight into the sampling of real networks.

Our findings are fourfold. First, *EcoNetGen* is useful in investigations of underlying topologies of interaction networks. Second, both the underlying pattern of species interactions and the strategy used to sample them had a large impact on the observed network structure. Third, sampling species according to the number of their interactions consistently resulted in more accurate estimates of the underlying network structure. Moreover, sampling according to module membership can produce good estimates of the structure within individual modules, but increases the risk of missing entire modules of species interactions altogether. Fourth and finally, we found that nested sets of interactions are easier to detect regardless of sampling strategy, potentially explaining the relative ubiquity of nestedness in empirical species interaction networks.

## Design and Implementation

### Generating networks

Networks with a specified number of modules and a variety of module topologies can be constructed with the software script *NetGen* in *EcoNetGen* (details and examples in Appendix S1-S2; scripts can be found in Appendix S5). We assume here that the complete networks we generate represent an ecological system encompassing multiple taxonomic groups, different habitats or types of interactions, here represented as network modules (Table 1). Therefore inter-module interactions are those that indirectly connect a taxonomic group with another (*e.g.*, an interaction between a bee and a plant that mostly interacts with hummingbirds), or interactions of a species that inhabits two habitat types over its lifetime (*e.g.* amphibians, who live part of their lifecycle in an aquatic subnetwork and the other part in a terrestrial one), or are species interactions involving species that connect different interaction modules (*e.g.* butterflies, which shift over ontogeny from herbivores to pollinators of the same plant).

*EcoNetGen* allows the construction of modular networks where the total network size (or total number of nodes, *N*), the average module size (*M*_*av*_), the average degree of the nodes (*k*), and topology of modules can be controlled. The simulated network can have either one or multiple modules, the sizes of which (*M*_*i*_) are drawn from an exponential distribution with average *M*_*av*_. Modules can have different topologies: random, scale-free, nested, bipartite nested, bipartite random, tripartite nested, and tripartite random (Appendix S1: Fig. A1). Networks may be uniform, such that all modules have similar structures (*e.g.* all scale-free) or may contain modules of various topologies (*e.g.* a combination of random and nested). When the generated network contains mixed modules, each module type is randomly chosen with given probabilities, so that only modules of certain types can be generated if the probability of the others is set to zero. Once the modules have been created, the links between nodes within each module can be changed, or rewired, with specified probabilities (*p*_*local*_) to randomize the initial structures. The nodes of the full network can then further be rewired (with probability *p*_*rew*_) to create connections among the modules.

We generated sets of modular networks with five different module topologies: random, scale-free, nested, bipartite nested, and mixed. Examples of adjacency matrices that correspond to the different module topologies are shown in Appendix S1. Although all network generation parameters can be specified by the user, we chose to fix the total network size at *N*=500, with average module size of 25, and average node degree of *k*=10 in order to reduce the number of parameters in our analyses. Though the average degree is fixed, the degree distribution can vary significantly depending on module type. Once the modules were constructed according to a given algorithm, nodes were randomly rewired to other nodes within the module with probability *p*_*local*_=0.1. This value was chosen to preserve the identity of the modules, but remove their “exact” algorithmic form. Nodes were further rewired to any node of the network with probability *p*_*rew*_=0.1, to create connections among modules.

### Sampling simulated networks

The motivation for examining different sampling designs applied to a full network is to explore how the most common practices used by a researcher with limited time or resources will affect the conclusions they draw about the underlying network structure. The sampling procedure, carried out in the script *NetSampler*, consists of picking *m* nodes that anchor the construction of the observed network, and then adding a number of first neighbor nodes (*nfn*) to each of these “anchoring” nodes (Appendix S3). Such a sampling design emulates a researcher studying a particular set of *m* species, and subsequently identifying those species that interact with the original set, as is often done when sampling animal-plant interactions (see Jordano 2016). The anchoring nodes and their neighbors can be chosen in different ways, as described below. We emphasize that only the observed interactions between nodes are included in the observed (or sampled) network. Therefore, two anchoring nodes that are connected in the original network will be connected in the sampled network only if one of the nodes is selected as a first neighbor of the other in the sampling process. In other words, an existing link between anchoring nodes is not automatically passed to the observed network.

### Sampling anchoring nodes

The anchoring nodes can be chosen according to different criteria: they can be chosen at random, according to the node’s degree (the number of interactions), according to abundances that are attributed to the nodes, or by attributing weights to each module such that species in one module (representing a particular interaction type or a taxonomic group) can be more or less likely to be chosen over species in other modules. Sampling procedures are implemented through *NetSampler*. Mathematical forms of these sampling distributions are available in Appendix S3, and are described below.

1. Random: *m* attempts to select nodes at random from the network are performed. The actual number of distinct anchoring nodes might turn out to be smaller than *m*, because the same node can be selected more than once. This sampling design represents a benchmark with which other sampling methods can be compared.
2. Degree of the node, *k*: the probability that a node is selected is proportional to its degree. The higher the degree of the node, the higher is the chance it will be included in the observed network. Again, *m* attempts are made, but fewer than *m* nodes might actually be included. The reasoning for such a sampling process is that a field biologist could choose to study a generalist species, whose interactions might be more likely to be observed because degree is sometimes correlated with abundance (Vázquez et al. 2009). Although in this sampling design we make no particular assumption about abundances, this relationship could be thought of as the underlying reason why interactions of species with higher degree may be more easily detected.
3. Abundances: an abundance value is attributed to each species (node) following three possible distributions: exponential, Fisher log-series, and lognormal (specified in *NetGen*, see equations B1, B6 and B8 in Appendix S2). Once the abundances have been attributed, *m* attempts to select nodes are made and the probability that a node is selected is proportional to its abundance. Abundances are attributed to each module independently and are not correlated to the degree of nodes or any network properties. This simulates a sampling process where the likelihood of sampling depends on abundances and favors the most abundant species of each module to be selected as anchoring nodes. This scheme differs from random sampling, where nodes have the same change of being selected irrespective of their module membership. Here each module is likely to have an anchoring node represented by its most abundant species. The process thus promotes uniform sampling of modules with random sampling within modules.
4. Module: sampling probabilities are assigned to the network modules, and within each module the probabilities associated with the nodes are uniform. In this way, species in some modules have a higher probability of being sampled than those in other modules, while the sampling probability is uniform within a given module. This incorporates the notion that some groups of species are easier to observe than others or that some researchers focus on particular types of interactions or taxonomic groups (see equation B9 in Appendix S2).

### Sampling interactions: choosing the neighbors of the anchoring nodes

Once the anchoring nodes have been selected, a subset of their interactions is sampled from the complete network to construct the observed network. Interactions are sampled in two ways: by specifying a maximum number of first neighbors (*nfn* is an integer and >1), or by specifying a fraction of the total number of neighbors per node (*nfn* < 1). Similar to the parameter *m*, *nfn* specifies the number of attempts to include neighbors: if a neighbor is selected twice, one attempt is lost. This is analogous to performing field observations for a limited time and observing several interactions between the same pair of species (Jordano 2016). If *nfn*=4, for example, a node with 2 links will very likely have its two neighbors included whereas a node with 8 links will have at most 4 of its neighbors included (with a range of 1-4 neighbors actually included). If *nfn*=0.5, on the other hand, the number of attempts per node is equal to half the number of its neighbors. Once the method for sampling interactions has been chosen, the actual neighboring nodes can then either be selected: (1) with uniform probability, or (2) with varying probabilities following an exponential distribution. In the latter case, these probabilities can be thought of as weights that represent interaction frequency, abundance, or a convolution of the two (Vázquez et al. 2005).

For each network, we sampled *m* anchoring nodes and randomly added *nfn* first neighbors for *m*=10, 20,…, 100 and *nfn*=5 or 10. This process simulates the sampling design used to build interaction networks from field data, where only a subset of species is repeatedly surveyed for their interactions. To demonstrate these methods, we performed 1000 replicates of sampling according to each scheme described above on each generated network.

### Network metrics

The number of components of the sampled network, together with the size distribution of these components, measure how well the between-module connectedness has been captured by the sampling procedure. Ideally a single component should emerge, matching the complete network. Therefore, for each sampling design, we calculated: (a) the size of the sampled network, *i.e.*, the total number of sampled nodes; (b) the number of components of the sampled network and; (c) the size of the largest component divided by the size of the sampled network, *i.e.*, the relative size of the largest component (RSLC). This last quantity measures the fraction of sampled network contained in its largest connected component.

Because we were interested in the overall topology of the network, we focused on metrics describing the size and number of components instead of assessing the internal structure of each component. Because the sizes of most components were typically small, measures such as average degree, clustering, average path length, or degree distribution would provide much information about the observed structures. However, since degree distribution is such a basic descriptor of networks, we explored how the degree of sampled nodes (in the observed network) correlate with their value in the complete network for different network topologies and sampling strategies.

## Results

We investigated how the incomplete sampling of large networks formed by several modular structures affected conclusions drawn about the underlying network structure, depending on the structure of the network, sampling intensity, sampling procedure, and the interaction between sampling procedure and network structure. Fig. 1 shows examples of bipartite networks generated with *NetGen* and the resulting sampled networks created by *NetSampler*. These results are for nested bipartite modules and *m*=50. The panels show the sampled nodes and links (red) embedded in the original network (blue) for three sampling methods (random, degree, and module). We note that sampling by module can leave entire modules hidden from the observer. This is the case of modules 2, 8, 9 and 14 in Fig. 1(e)-(f). Analogous results for a network with mixed modules are shown in Appendix S4.

**Figure 1.**
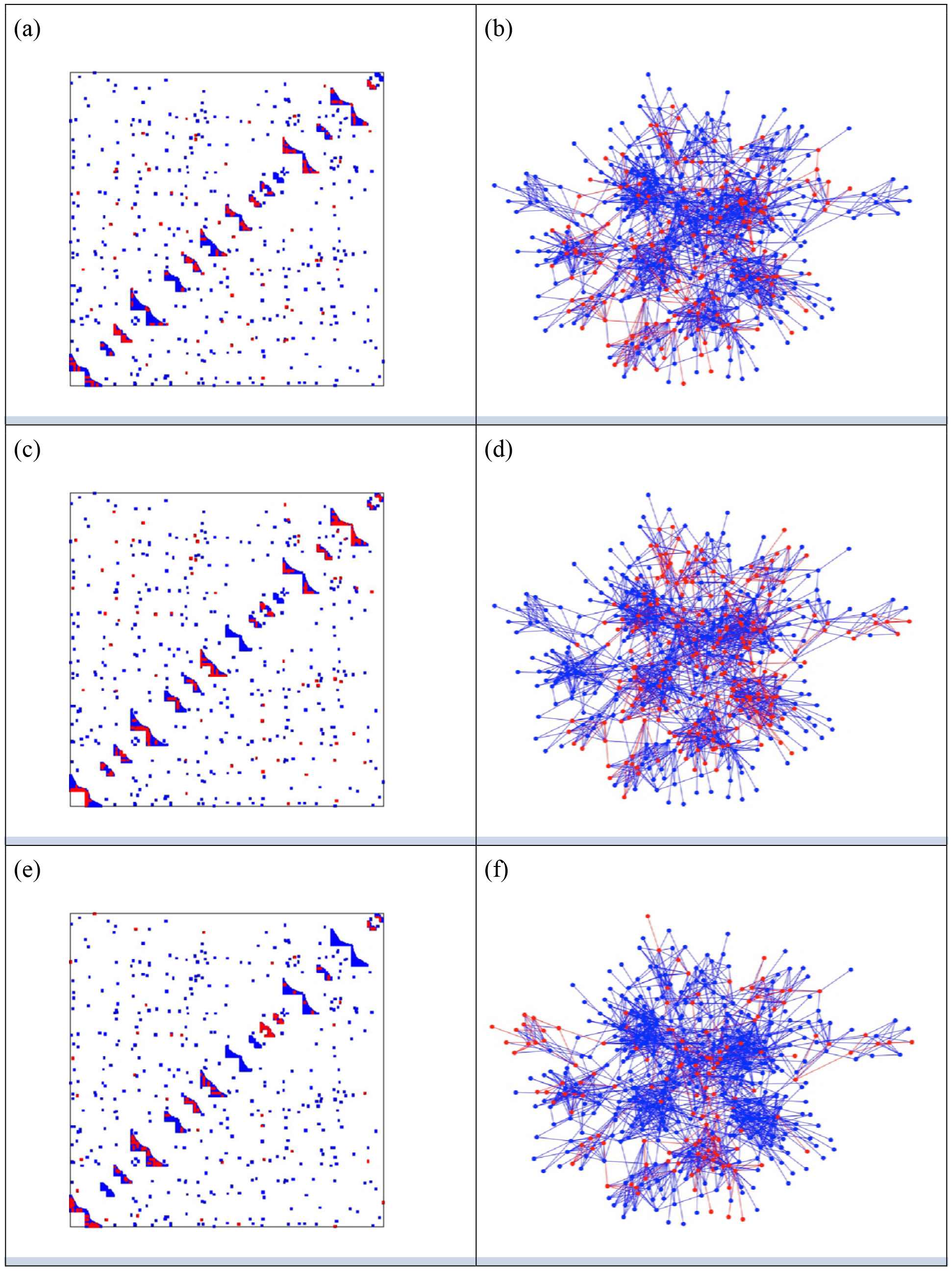
Adjacency matrices and network structure for a network with bipartite nested modules. Sampling occurred on *m*=50 anchoring nodes and adding up to 10 first neighbors. The complete network has 16 modules with sizes 49, 21, 27, 55, 31, 25, 41, 38, 17, 21, 15, 55, 12, 58, 12 and 13. The average degree is 7.5 and average module size is 31.25. Key nodes were chosen randomly (a)-(b), according to degree (c)-(d) or to module preference (e)-(f). Nodes and links in red represent the sampled species and interactions in each case. The number of connected components in each case is 12, 6 and 13 respectively.

Clustering of sampled nodes are apparent and visually different between sampling methods, which can be analyzed according to network metrics described in *Methods*. Relative size of the largest component varies with underlying network topology and sampling method (Fig. 2-3, with RLSC as a function of *m* for 5 sampling methods for bipartite nested modular networks and mixed module networks, respectively). Interactions between underlying network structure and sampling design were quantified by number of components, RLSC, and size of the sampled network (Fig. 4). The modular structure of the complete network led to sampled networks consisting of several disconnected components corresponding to nodes from a single module or from a small group of modules. In Fig. 4, for example (*N*=500, *m*=50, and *nfn*=5), sampled networks typically had 12 disconnected components comprising approximately 150 sampled nodes.

**Figure 2.**
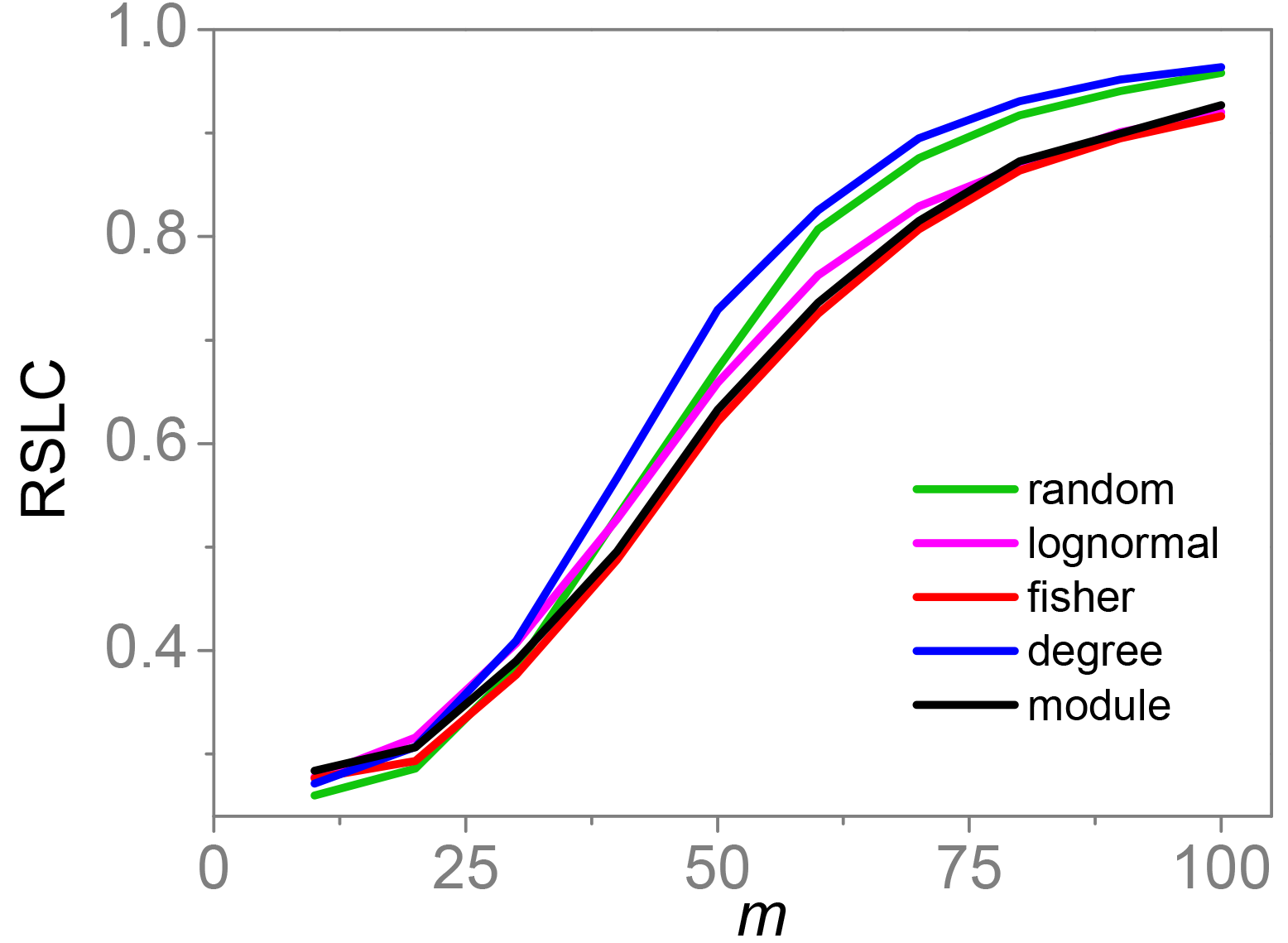
Relative size of the largest component (RSLC) is shown for a network with bipartite nested modules as a function of *m* for *nfn*=10, with line color indicating sampling design.

**Figure 3.**
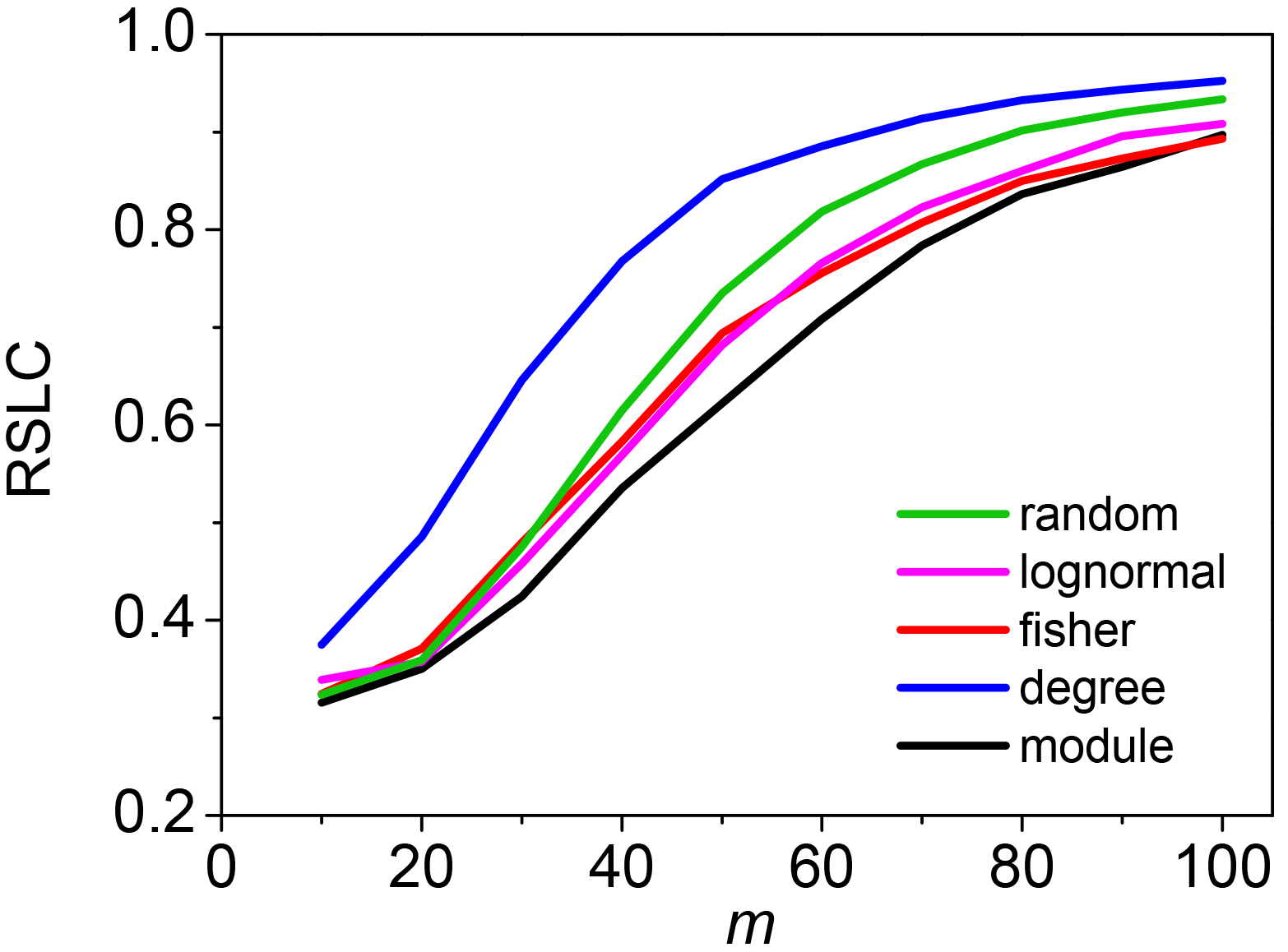
The relative size of the largest component (RSLC) is shown for a network with mixed modules as a function of *m* for *nfn*=10, with line color indicating sampling design.

**Figure 4.**
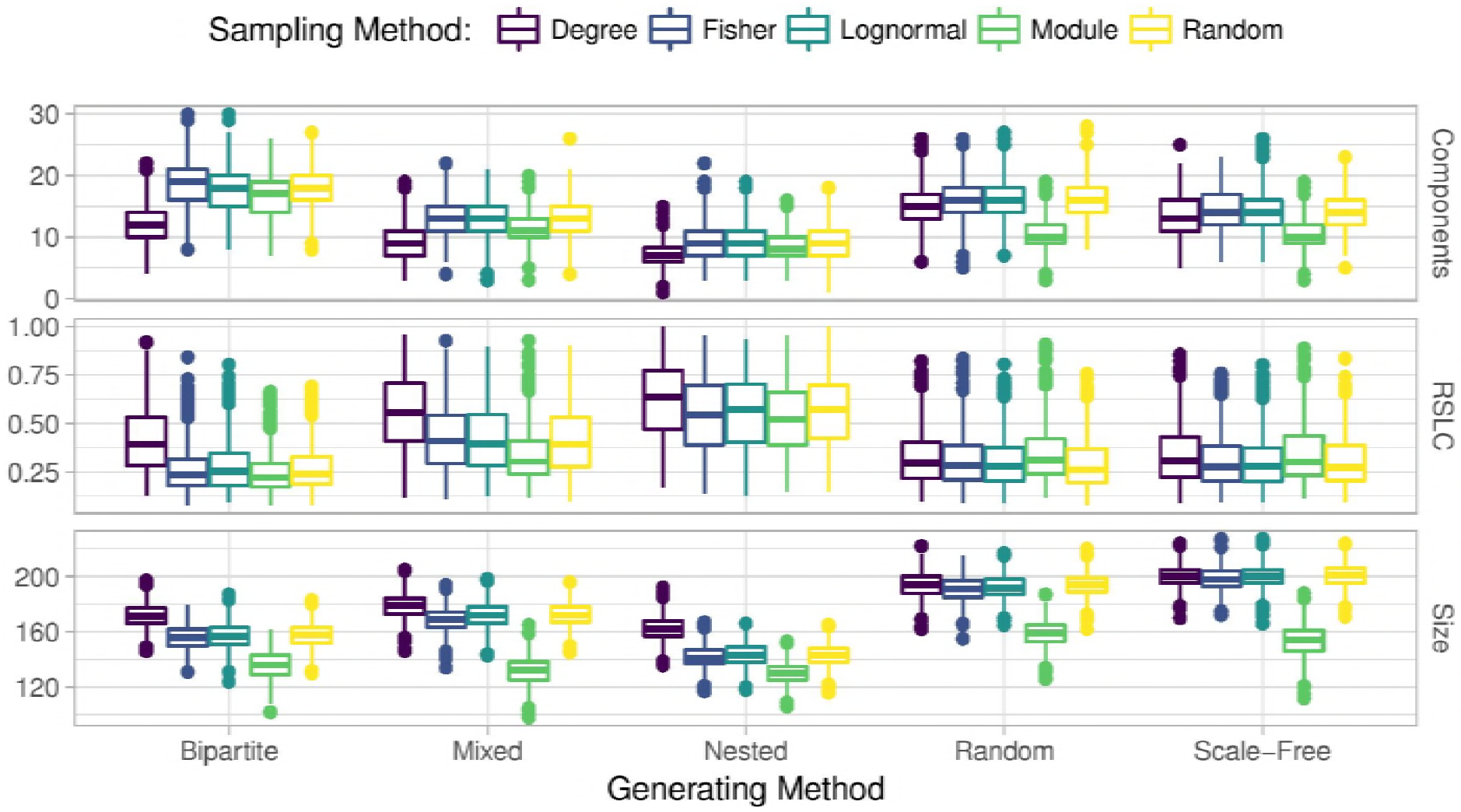
Boxplots depicting average and standard error for number of connected components, relative size of largest components (RLSC), and size of sampled network (measured by number of nodes) for generated networks with *N*=500, *m*=50 and *nfn*=5, sampled with multiple methods. For each method, the generated network was sampled with 1000 replicates.

We also found that the degree of sampled key nodes correlated with their *true* degree in the complete network (Fig. 5). We demonstrate this for the bipartite network with *m*=50, *nfn*=10 and three sampling methods. In all cases, there is a clear correlation between the true degree (corresponding to the complete network) and sampled degrees, represented by the straight ridge on the contour plots. Many of the patterns network ecologists are interested in are related to the degree of generalization or specialization of species, so even if the “true” degree cannot be determined by sampling, this demonstrates that all sampling schemes can capture reliable estimates of the degree of a species relative to others at low degrees of connectedness. Sampling by degree is one way to mimic a passive sampling where species with more interactions are more likely to be sampled. Only sampling by degree captures a saturation of the node degree, which is imposed here by the number of first neighbors added to the key nodes, *nfn*=10.

**Figure 5.**
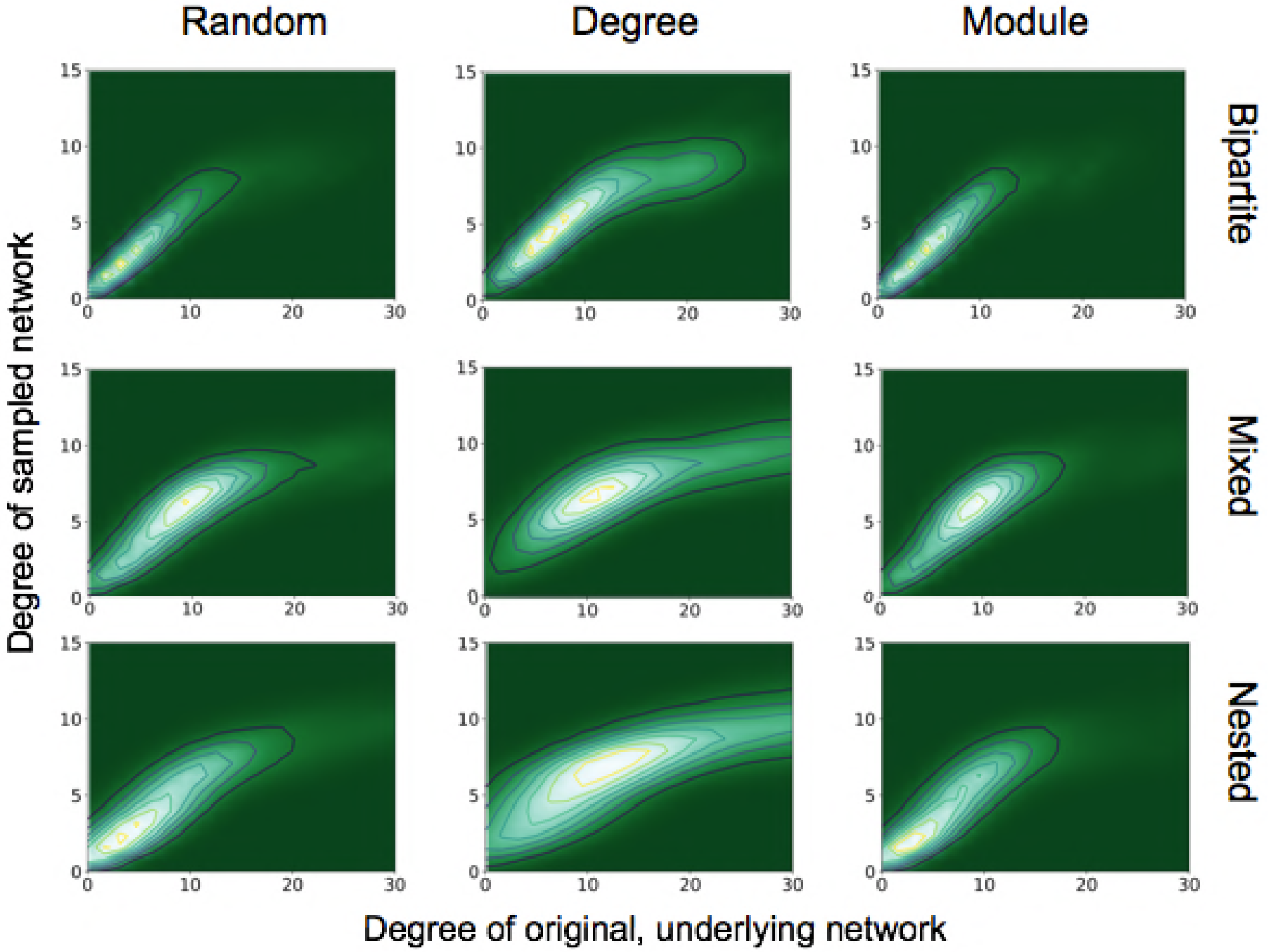
Degree of the sampled network versus degree of the complete network for bipartite, mixed and nested networks with *m*=50 and *nfn*=10. Only the sampled key nodes are shown. From left to right the sampling methods are: random, degree and module. Contour plots were generated with 50 realizations of samplings on each network.

Sampling by degree, by definition, tends to find more anchoring nodes at the center of the modules and few in the periphery. Random sampling, on the other hand, picks relatively more peripheral nodes, as can be seen by comparing Fig. 1(b) with 1(d) and 2(b) with 2(d). These general features are also present in nested, scale-free, and random networks (not shown). The aggregate statistics of the sampled network properties we calculated are summarized in Fig. 4 for networks with five different module types and five sampling methods. Of the major metrics we investigated, we observed the following trends:

1. Number of connected components: Networks with nested modules have the smallest number of connected components and show the highest level of between-module connectedness. The other network types did not show significant variation in the number of connected components across the different sampling designs.
2. Relative size of largest connected component: Networks with nested modules had largest sampled components containing up to 60% of the entire number of observed nodes, whereas the largest component of the other network types represents only 30% of the observed nodes, revealing a much lower degree of connectedness between modules than in nested networks. An exception is the mixed network sampled by degree, whose largest component contained 56% of the nodes of the observed network.
3. Size of sampled network: Networks with nested and bipartite nested modules always produced the smallest sampled networks, independent of the sampling procedure (Fig. 4). Networks with scale-free modules, on the other hand, produced the largest observed networks, followed closely by those with random modules. On average, observed networks that were sampled from full networks with nested modules were 72% smaller than those sampled from networks with scale-free modules. This is because species with high degree in the scale free module will be in hubs that are also connected with species in other modules. Hence, once such species are sampled, we are likely to sample several other nodes from other modules. Mixed networks fell in between these two cases, as expected.
4. Degree of sampled nodes: All sampling methods show linear correlation between the *true* and sampled degrees for key nodes. Sampling by degree goes beyond the linear correlation and captures the saturation at degree *nfn* imposed by the number of first neighbors added to the key nodes. No other sampling method generated this outcome.

## Discussion

Although complete ecological networks found in nature may be incredibly large, efforts to understand the structure or dynamics of these empirical systems have focused on smaller, tractable subcomponents of the actual networks due to limitations of time, energy, and budget (Burkle & Alarcón 2011). Moreover, most field-based attempts to quantify ecological networks limit the types of interactions being measured to particular species of interest. In our formalization, individual studies would generally be examining a single module that exists within a larger universe of interactions, defined here as the complete or underlying network. For example, a plant-pollinator module is depicted as a tightly interconnected bipartite network (Bascompte & Jordano 2007), where the trophic interactions of its constituent species are part of a separate trophic network that may be ignored.

The complete network, encompassing different types of interactions and taxonomic groups, as well as the structural heterogeneity depicted among its subnetworks (here represented by the distinct modules), is rarely addressed. Nonetheless, the ecological and evolutionary dynamics of these modules in the community are hardly independent of each other. Modules may emerge naturally due to the sparseness of interactions across space, time, or even as the result of coevolutionary forces (Olesen et al. 2007; Beckett & Hywel 2013; Andreazzi et al. 2017). Still, the effects of interactions in one module can propagate across the system (McCann et al. 2005; Rooney, McCann et al. 2006). The groups of species and types of interactions one targets when conducting fieldwork will define the type and size of the network studied. Analytical techniques that simultaneously address multiple types of interactions and ecological outcomes (*i.e.* multilayered networks; Pilosof et al. 2017; Genrich et al. 2016) will require understanding the bias imparted by sampling strategies in order to deal with the insurmountable diversity of organisms and interactions in real communities. Thus, if we desire to understand the relationships between structure and function, we should ultimately aim to obtain the most accurate depiction of the structure encompassing all elements potentially affecting function.

We attempted to quantify how much of the idealized network is observable, and what systematic biases may exist as a function of both the designs used to sample species interactions and the complete network structure. To this end, we provide the software *EcoNetGen*, which can be used to explore many other features of biases introduced by sampling methods. We investigated only a few of these features, related to the between-module connectedness of the observed network. We highlight that the scripts contained in *EcoNetGen*, *NetGen* and *NetSampler*, may be used to form null models and simulated networks in other studies.

Our analyses using *EcoNetGen* point to four main results. First, sampling design has a large impact on the properties of the observed network. Sampling according to species degree seems to be the only method that consistently generates nearly complete networks, as it produces the largest and more connected observed networks, with the smallest number of components. For most ecological systems, natural history studies can provide intuition about which species are most likely to be the highly-connected in the network, and which species are highly specialized. Moreover, it is relatively easy to identify the highly-connected species through incomplete sampling schemes (Pires et al. 2017). Thus, a combination of natural history information and sampling schemes focused on highly-connected species may provide the most accurate description of ecological networks.

Further, networks comprised of bipartite nested and nested modules generally result in poorly sampled networks that are small and have several disconnected components. This suggests that networks with these types of structures demand greater sampling effort than networks with random or scale-free modules, for instance. Although real networks are in-between these different structural patterns explored here, this highlights that special care should be taken with sampling design, since the interplay between sampling and structure will affect how the representativeness of the sampled network. The positive message is that sampling from networks with bipartite nested modules results in observed networks with a large number of small components, meaning that each module is well sampled. The challenge to build more representative networks is to devise ways to sample the connections among modules, which are rarely observed by the different sampling schemes for these types of networks. In comparing sampling designs, it is clear that sampling by module produces by far the smallest observed networks for all topologies (Fig. 4). Sampling by degree, on the other hand, produces the largest sampled networks (as measured by the relative size of the largest component) and is therefore most representative of the underlying complete network. For networks with random and scale-free modules, sampling by degree produced similar results compared to random sampling, but for nested and bipartite nested networks, sampling by degree always produced significantly larger observed networks than random sampling. Interestingly, sampling by abundance does not seem to be appropriate for nested or bipartite nested networks, because the results are only slightly better than sampling by module. For random and scale-free networks, sampling by abundance produces observed networks that are only slightly smaller than those produced by sampling by degree.

Second, nested modules are better represented in sampled networks than other module structures. Since the renewal of the interest in ecological networks in recent decades, nestedness has played a central role in the literature and has been reported in a wide variety of systems described as bipartite networks (e.g. Bascompte et al. 2003; Guimarães et al. 2006; Joppa et al. 2010). However, the relevance of nestedness has been contested (Staniczenko et al. 2013; James et al. 2012), and mechanisms such as abundance heterogeneity and sampling have been invoked as underlying causes of the pervasiveness of the nested pattern (Vázquez et al. 2009). The fact that nested modules are overrepresented compared to other module types in sampled networks suggests that a major underlying reason for the ubiquity of nestedness in empirical networks is that sampling strategies employed in empirical studies are successful in thoroughly sampling nested subnetworks (Nielsen & Bascompte 2007), but may not perform as well when sampling non-nested subnetworks. Similarly, networks with unipartite nested modules (Cantor et al. 2017) stand out as providing observed networks with the most closely connected of all topologies. Sampling by degree, *i.e.*, with focus on those species likely to establish more interactions, is therefore the recommended procedure for sampling networks with mixed modules, but it may overestimate the relative frequency of nested modules because non-nested modules are harder to thoroughly sample. Testing additional sampling designs capable of identifying other structures will aid in understanding the relative frequency of the different structural patterns in real networks.

Third, the size of the observed network does not depend significantly on the sampling method, but depends strongly on the underlying network topology. As shown in Fig. 1, the size of the network at *m*=50 is smaller for nested and bipartite nested networks, independent of the sampling criterion, whereas networks with scale-free modules produce the largest sampled networks. This happens because the degree distribution in nested networks is very heterogeneous and more anchoring nodes will likely have fewer than *nfn* neighbors. Sampling according to module always produces small observed networks, but network size does not change much for the other methods. This suggests that sampling the entire network will be difficult no matter which sampling strategy is chosen, and perhaps the best strategy is one of iterative sampling, where the structure of a partially sampled network is analyzed and sampling is resumed using the sampling design that best suits the uncovered structure.

Fourth and finally, sampling according to module generally results in small observed networks with a small number of observed components. This means that the method may detect most of the inner structure of some modules, but does thoroughly sample a part of the network, allowing for identification of interactions in multiple modules. This type of sampling is arguably the most pervasive in the network literature where a certain type of interaction or taxonomic group is exhaustively sampled. Our simulations show that sampling by module may give a thorough depiction of the module but may also point to other modules, which can be then sampled according to the most adequate sampling design.

Because properties of the network that determine the most efficient sampling strategy (such as species degree and module topology) are not fully known prior to undertaking a study, we recommend an iterative sampling approach, where the strategy can be adjusted as network properties are revealed. In recent years, interest in ecological network theory has grown exponentially (Costa et al. 2007; Borrett et al. 2014; Delmas et al. *in review*) while our understanding of empirical systems has lagged behind, in part due to the difficult and time-intensive nature of field data collection and sample processing. Only by integrating a formal understanding of how empirical efforts reflect or bias estimation of the underlying network of species interactions can we confront theoretical models with our observations of natural systems.

## Acknowledgements

This work was conducted as a part of the Ecological Network Dynamics Working Group at the National Institute for Mathematical and Biological Synthesis, sponsored by the National Science Foundation through NSF Award #DBI-1300426, with additional support from The University of Tennessee, Knoxville. MAMA was partly supported by FAPESP (grants #2016/06054-3 and # 2016/01343-7) and CNPq (grant #302049/2015-0).

## Supplementary Material

**Appendix S1:** Generating networks with *NetGen*

**Appendix S2:** The abundance distributions

**Appendix S3**: Sampling simulated networks with *NetSampler*: an example

**Appendix S4:** Adjacency matrices and network structure for a network with mixed modules

**Appendix S5:** *NetGen* and *NetSampler* Python and Fortran scripts

